# Mating-induced ecdysone in the testis disrupts soma-germline contacts and stem cell cytokinesis

**DOI:** 10.1101/2023.10.16.562562

**Authors:** Tiffany V. Roach, Kari F. Lenhart

## Abstract

Germline maintenance relies on adult stem cells to continually replenish lost gametes over a lifetime and respond to external cues altering the demands on the tissue. Mating worsens germline homeostasis over time, yet a negative impact on stem cell behavior has not been explored. Using extended live imaging of the *Drosophila* testis stem cell niche, we find that short periods of mating in young males disrupts cytokinesis in germline stem cells (GSCs). This defect leads to failure of abscission, preventing release of differentiating cells from the niche. We find that GSC abscission failure is caused by increased ecdysone hormone signaling induced upon mating, which leads to disrupted somatic encystment of the germline. Abscission failure is rescued by isolating males from females but recurs with resumption of mating. Importantly, reiterative mating also leads to increased GSC loss, requiring increased restoration of stem cells via symmetric renewal and de-differentiation. Together, these results suggest a model whereby acute mating results in hormonal changes that negatively impact GSC cytokinesis but preserves the stem cell population.

**Graphical Abstract:** 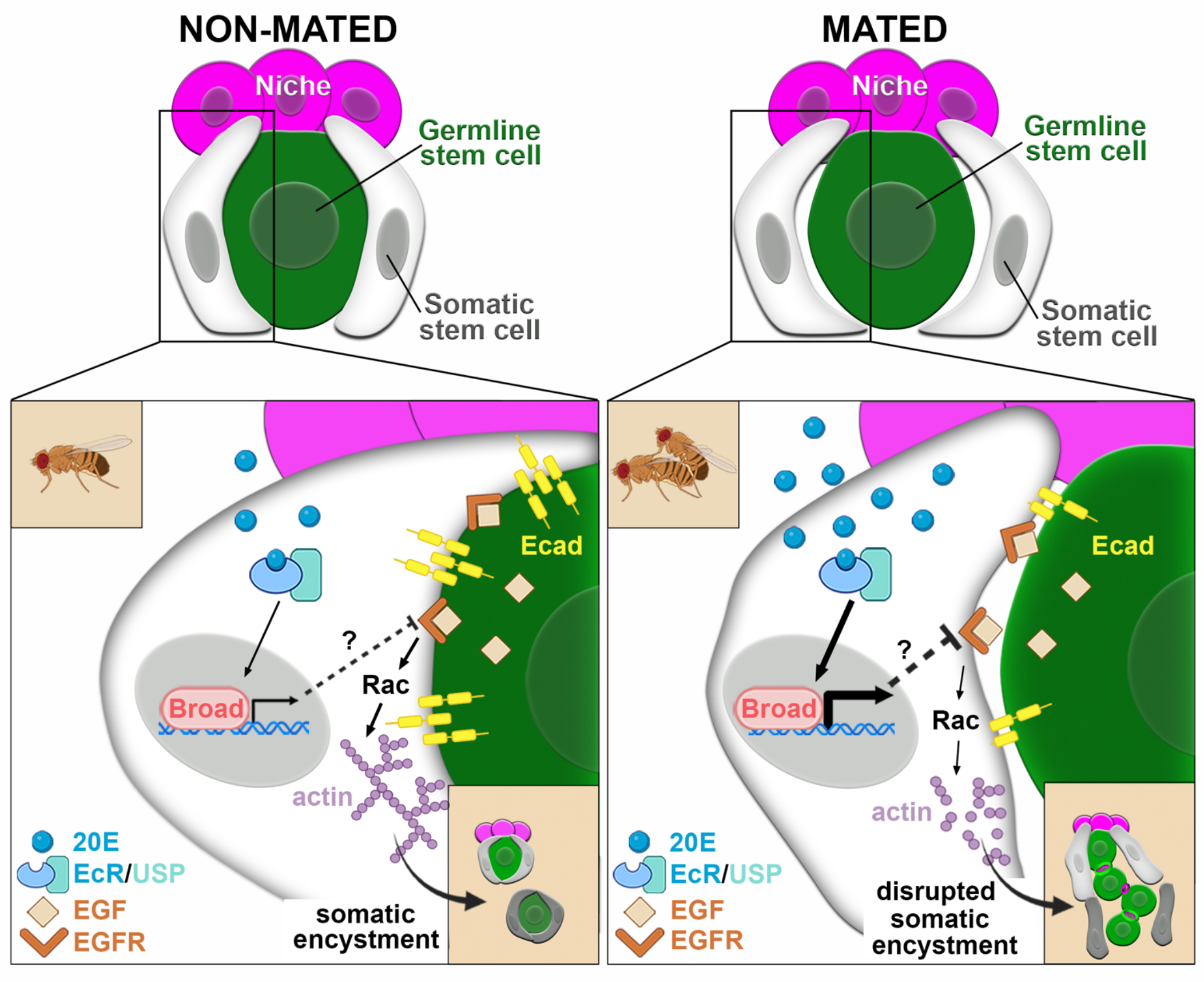

## Introduction

Tissue homeostasis is accomplished by balancing stem cell self-renewal and production of differentiating daughter cells. Tissues engage multiple mechanisms to uphold this balance under stressful conditions.^1^ Mating is a natural physiological process but can be detrimental to tissue homeostasis.^2^ Increased production of sperm or eggs depends on the ability of stem cells to respond to and compensate for demand. Studies have shown that long-term mating combined with aging or nutrient deprivation results in loss of stem cells or over-proliferation of undifferentiated cells.^3,4^ However, much less work has been done to identify the proximate effects of mating stress, alone, on stem cell biology. While some changes in stem cell behaviors can be identified from fixed tissue^5,6^, these analyses are limited in the ability to resolve acute changes in cellular dynamics.

The *Drosophila* testis niche is an ideal model system to address how physiological changes directly impact stem cell behavior. The testis contains two resident stem cell populations: germline stem cells (GSCs) and somatic cyst stem cells (CySCs). As in most animals, there is an obligatory association between soma and germline.^7^ In the fly testis, germ cell differentiation requires full encapsulation of each GSC daughter by two somatic cyst cells of the CySC lineage in a process called encystment (Fig.1A).^8,9^ Soma-germline contacts are then stabilized and maintained by adherens and septate junctions as they co-differentiate.^10,11^ Using extended live imaging of the stem cell niche, we find that mating disrupts GSC cytokinesis by decreasing soma-germline contacts. Abscission failure caused by defective encystment is mediated by increased ecdysone signaling upon mating, is significantly reduced when mating stress is removed, and recurs when mating is reintroduced, demonstrating a consequential impact on germline homeostasis.

**Fig. 1.**
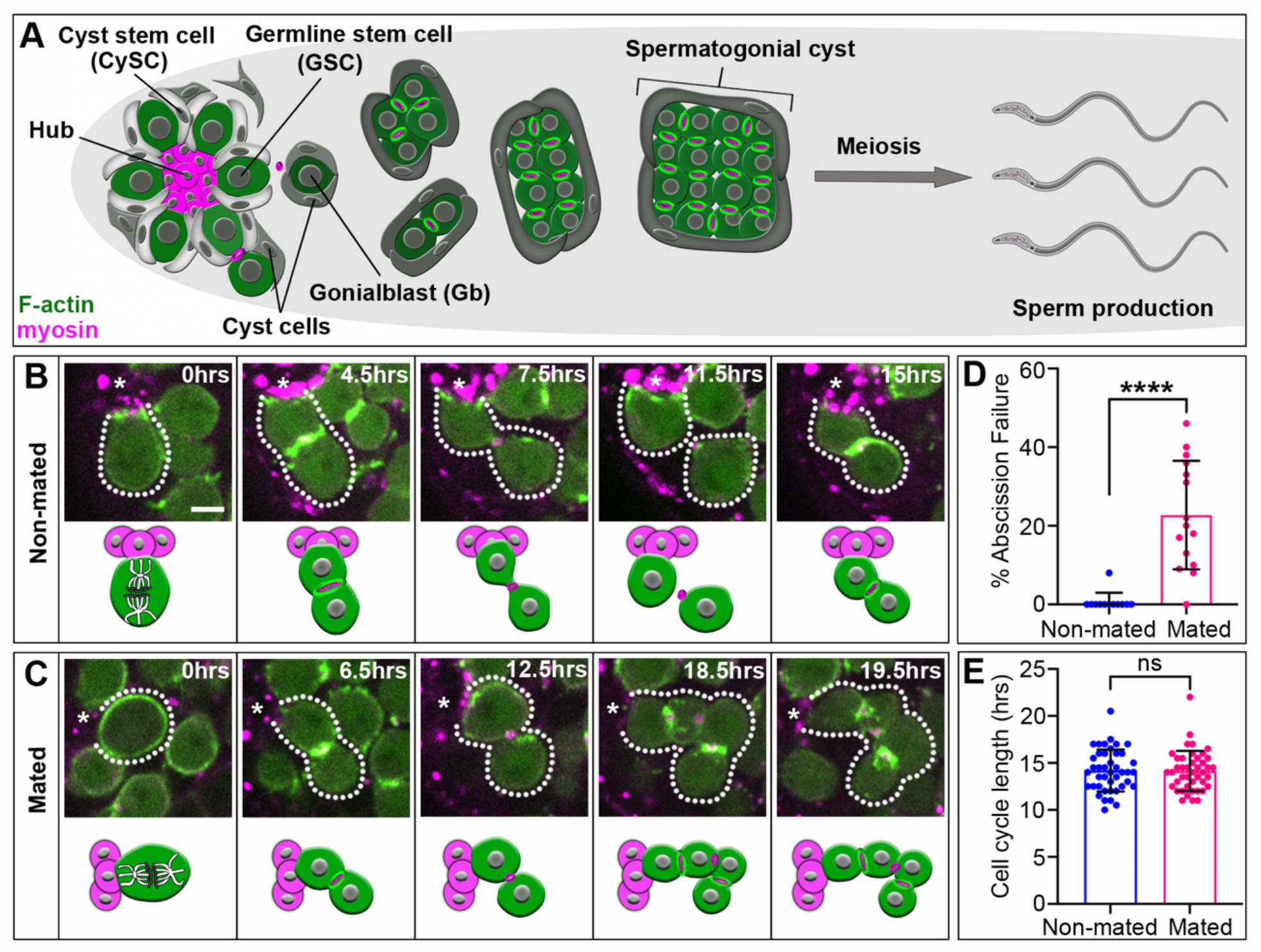
Mating causes a significant increase in GSC abscission failure. (A) Diagram of testis stem cell niche and germline development. (B and C) Time-lapse of nanos-ABD-moe::GFP and Myo-mCherry (sqh-sqh::mCherry) during GSC divisions in (B) non-mated and (C) mated animals. Each image is 1-4 z slices to capture the division plane. Asterisks: hub; hours (hrs); scale bar: 5µm (D) Percent GSC abscission failure under non-mating (n=1/135 divisions in 12 testes) and mating conditions (n=40/167 divisions in 15 testes). \*\*\*\**P*<0.0001. (E) GSC cell cycle length in non-mated (n=42 GSCs in 10 testes) and mated animals (n=43 GSCs in 14 testes). ns, not significant.

## Results and Discussion

### Mating disrupts the release of GSC daughter cells from the niche

Mating has been shown to induce increased GSC numbers and cycling rates, but these changes do not fully explain how mating contributes to tissue degeneration or tumorigenesis.^4– 6^ Aging stress causes early defects in GSC cytokinesis eventually leading to reduced numbers of differentiating germ cells—a hallmark of diminished homeostasis.^12^ To address whether mating impacts GSC cytokinesis, we performed extended live imaging^12,13^ to directly visualize GSC divisions and cytokinesis in testes of males that underwent 2 days of mating in a 3:1 virgin female to male ratio. In testes from non-mated males, GSCs execute a reproducible cytokinetic program that differs from typical cytokinesis observed in most other cells.^13^ Following contractile ring disassembly in mitosis, a secondary F-actin ring forms at the intercellular bridge between the GSC and daughter cell (Fig1.B). This F-actin ring then prevents cytokinesis progression until its disassembly, at which point cytokinesis resumes and completes with the final severing of membranes between daughter cells known as abscission. Under non-stress conditions, GSC abscission is extremely robust and nearly always completes prior to the next mitotic division (Fig.1D). By contrast, we found that GSCs in testes from mated males proceed normally through cytokinesis but exhibit significant abscission failure (Fig.1C-D, p<0.0001). This results in a chain of connected daughter cells remaining attached to the niche (Fig.1C), preventing release of differentiating germ cells. Thus, despite increased need for germ cells upon mating our data suggests the immediate effect on GSCs decreases the release of daughters from the niche.

Previous studies analyzed mitotic or synthesis (S) phase indices on fixed tissues and found that even short-term mating can induce a faster cycling rate in both male and female GSCs.^5,6^ As we previously showed that fast cycling can lead to abscission failure^12^, we first asked if changes in cycling rate cause the cytokinetic defects in GSCs upon mating. Our extended imaging (24 hours) allowed us to directly quantify cell cycle lengths of individual GSCs.

Consistent with existing literature, we observed variability in cell cycling length (10-22hrs) with an average cycling rate of 14hrs (Fig.1.D).^12-14^ However, we find that GSCs from mated males have both similar variability and identical average cycling rates compared to non-mated males (Fig.1.D, p=0.8418). To confirm these results, we determined the S phase index of GSCs in fixed tissue by exposing testes to 5-ethynyl-20-deoxyuridine (EdU) prior to fixation. Again, we found no significant difference in GSC S phase index between non-mated and mated testes (Fig.S1, p=0.9102). Together, these data suggest that mating-induced abscission failure is not caused by increased cycling rates in GSCs.

### Mating causes defects in somatic encystment of germ cells

We previously identified a critical role for somatic encystment in the completion of GSC cytokinesis through abscission.^13^ Importantly, GSC abscission failure caused by disrupted encystment occurs in the absence of changes to GSC cycling rates (Lenhart lab unpublished). As this is precisely the phenotype in mated flies (Fig.1C), we investigated whether soma-germline interactions are disrupted upon mating. Current analyses of encystment rely on visualizing large morphological changes or full-tissue quantification of adhesion proteins.

However, our previous work indicates that testes with subtle encystment disruptions can appear wildtype in morphology while having significant effects on GSC abscission.^13^ Therefore, we established a method for detecting subtle disruptions in soma-germline contacts focusing on 2-cell cysts, as encystment is complete by this stage (Fig. 2A). We quantified the mean fluorescence intensity of the adherens junction protein E-cadherin (E-cad) specifically at soma-germline interfaces and determined the fold change in fluorescence relative to controls (Fig. 2A-E). First, we validated this method in testes with soma expression of a dominant negative form of Rac1-GTPase (Tj>DN-Rac1; F.2C), previously shown to subtly disrupt encystment yet induce GSC abscission failure ^13^, and observed significantly reduced E-cad intensities at soma-germline junctions compared to controls (Fig.2B-C,F). Importantly, quantification of E-cad in non-mated versus mated males also revealed a significant decrease in adherens junction formation between soma and germline (Fig.2D-F p<0.0001), suggesting that somatic encystment becomes disrupted upon mating.

**Figure 2.**
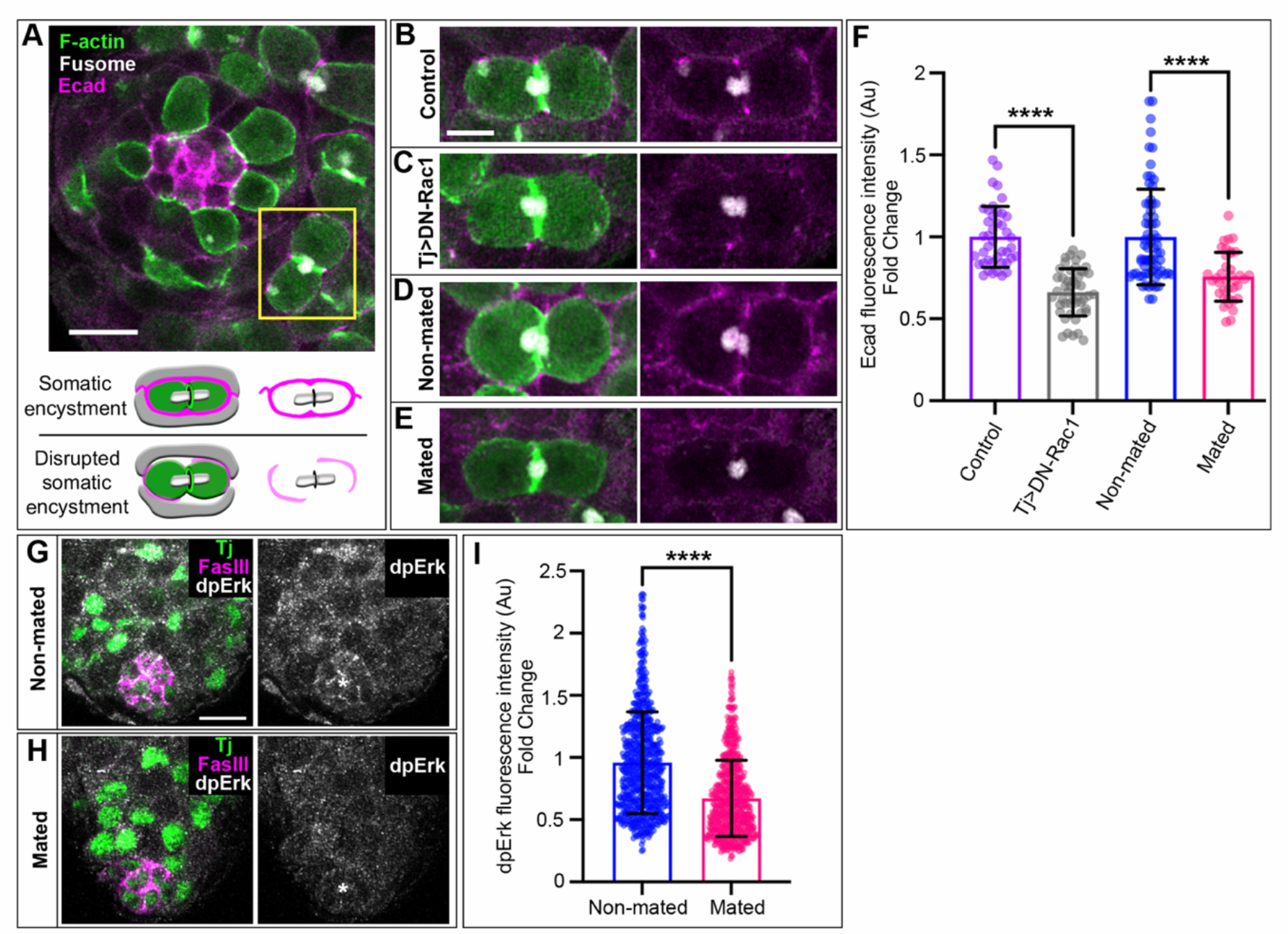
Mating leads to decreased adhesion and EGFR signaling between soma and germline. (A) Apical tip of the testis showing 2-cell cyst (yellow box) and diagram of normal versus disrupted somatic encystment. Scale bar: 10µm (B-E). Magnified view of 2-cell cysts from (B) a control testis with typical E-cad between soma-germline, (C) a testis with somatic expression of Rac1^DN^ and diminished E-cad, (D) a non-mated testis and (E) a mated testis showing decreased E-cad between soma and germline. Scale bar: 5µm (F). Quantification of E-cad fluorescence intensity represented as fold change relative to the respective controls. (G,H) Immunofluorescent staining of somatic nuclei (green, (Tj)), niche cell junctions (magenta (FasciclinIII/FasIII)) and dpErk (gray) in a (G) non-mated testis and (H) a mated testis with diminished dpErk staining. scale bar: 10µm (I) Quantification of dpErk fluorescence intensity in somatic nuclei represented as fold change \*\*\*\**P*<0.0001.

Encystment is mediated by epidermal growth factor receptor (EGFR) activation on somatic cell membranes, which triggers two intracellular pathways: 1. Activation of the Rac GEF, Vav, which enables rearrangement of the actin cytoskeleton to encapsulate germ cells and 2. Transcriptional changes through phosphorylation of Erk/MAPK.^8,9^ The fluorescence intensity of dpErk immunostaining can be used as a readout for the level of EGFR activation which impacts both the transcriptional and encystment pathways.^11^ We found a significant decrease in somatic nuclear dpErk in mated males compared to non-mated males (Fig.2G-I), suggesting decreased EGFR pathway activation. As EGF signaling is the critical pathway controlling encystment, these data along with the direct assessment of soma-germline junctions strongly suggest that mating disrupts somatic encystment of the germline, causing GSC abscission failure.

### Germ cell depletion does not cause GSC abscission failure

Previous work has shown that changes in GSC biology with mating are induced by the act of mating itself but not by pheromone sensing or exposure to courtship behavior.^6^ However, which physiological effect of mating is responsible for altered stem cell behavior remains unclear. At the tissue level, the most critical consequence of mating is depletion of the differentiating germ cell population, a situation which could feed back to change GSC behavior. Therefore, we first tested whether depletion of sperm in the absence of mating might be sensed by GSCs leading to abscission failure. We triggered ejaculation by optogenetically activating Crz neurons expressing red-shifted channelrhodopsin CsChrimson in a pattern mimicking the number of mating events a fly would experience in our paradigm thereby forcing depletion of sperm in the absence of mating (Fig.3A’).^15^ Interestingly, we did not observe significant GSC abscission failure upon sperm depletion (Fig.3C). We next tested if a more severe depletion of differentiated germ cells in the absence of mating was sufficient to induce GSC abscission failure. By expressing an apoptotic factor, Hid, in differentiating germ cells via Bam-Gal4, we can retain the niche and resident stem cells, but effectively kill transit amplifying germ cells, depleting the testis of all germ cells past the 2-cell cyst stage (Fig.3B’). Surprisingly, we find that catastrophic loss of differentiating germ cells did not cause abscission failure (Fig.3C). Together, this data suggests that depletion of germ cells does not cause GSC abscission failure and that a different aspect of mating is leading to defects in the stem cell pool.

**Figure 3.**
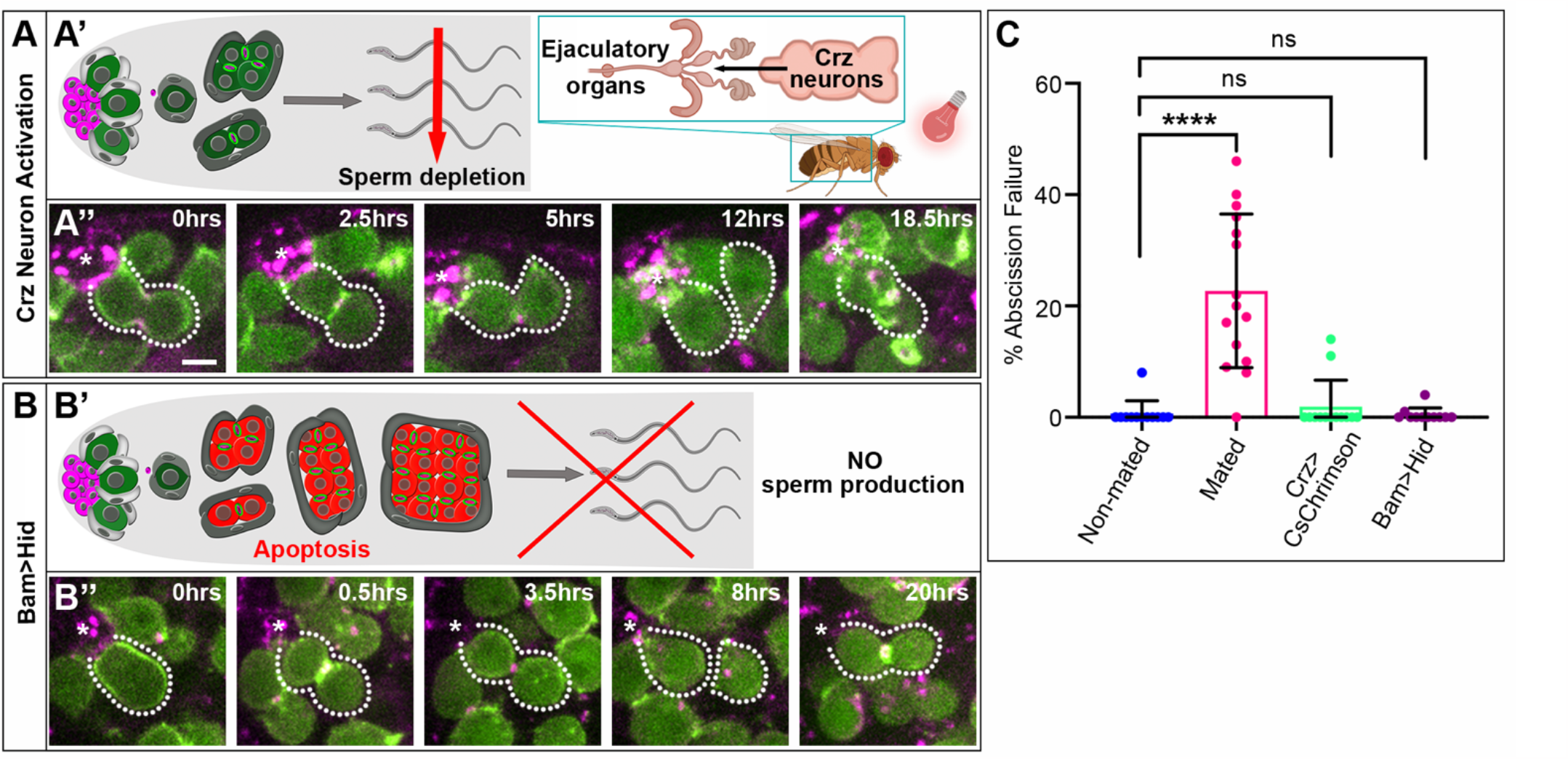
Sperm or germ cell depletion does not cause GSC abscission failure. (A’’-B’’) Time-lapse of nanos-ABD-moe::GFP and Myo-mCherry (sqh-sqh::mCherry) during GSC divisions in testes of (A’’) males with optogenetic activation of Crz neurons and (B’’) males in which germ cells from the 2-cell cyst onward express the apoptotic factor Hid to induce cell death. (A’ and B’) Diagrams of how sperm or germ cells were depleted in the absence of mating. (D) Quantification of GSC abscission failure (non-mated and mated data repeated from Fig.1).

### Ecdysone signaling is induced by mating and is sufficient to disrupt encystment, causing GSC abscission failure

Ecdysone signaling increases in the female gonad following mating causing increased GSCs divisions.^5^ Although male accessory organs have increased biosynthesis of ecdysone during mating^16^, whether released ecdysone affects the testis or male GSC function is unknown.

Altered ecdysone signaling in the female gonad has been shown to perturb the formation of cytoplasmic extensions of somatic escort cells to early germ cells^17^, a process analogous to somatic encystment in the testis. Furthermore, ecdysone and EGFR signaling have been shown to act antagonistically in male somatic cells^18^; in contrast to EGFR, ecdysone is suggested to promote activation of Rho^19^, which antagonizes Rac activity that in turn is essential for encystment.^8^ Thus, we investigated whether mating increases ecdysone signaling in testis soma leading to disrupted somatic encystment and subsequent GSC abscission failure.

To assay ecdysone activity, we quantified fluorescence intensity of the known ecdysone target, Broad-core.^20^ We identified somatic nuclei by Traffic Jam staining and quantified Broad levels between non-mated and mated testes (Fig.4A-B). We found that mating induced a significant increase in Broad accumulation (Fig.4C), suggesting that ecdysone signaling in soma is indeed increased upon mating. Next, to address if this increased activity is the proximate cause of GSC abscission failure upon mating, we fed males the steroid hormone 20-hydroxyecdysone (20E) to induce signaling in the absence of mating.^20,21^ Flies were fed a mixture of apple juice and blue dye supplemented with either vehicle control (1% ethanol, lacking 20E) or 20E at a concentration of 1mM on filter paper each day for 2 days to closely mimic the mating-induced increases in ecdysone signaling. Excitingly, 20E feeding in the absence of mating was sufficient to induce a significant increase in GSC abscission failure (Fig.4D-F). Abscission failure with mating is caused by diminished somatic encystment. Thus, we next questioned whether the abscission failure induced by 20E feeding was also caused by decreased soma-germline contacts. Quantification of E-cad after 20E feeding revealed significantly decreased levels at soma-germline contacts (Fig.4G), suggesting that increased ecdysone signaling is the proximate cause of encystment disruptions and GSC abscission failure upon mating. This result is particularly striking as 20E feeding is *not* sufficient to induce increased GSC cycling in females despite the essential role of ecdysone in that process.^5^ Future work will be directed toward understanding the differing mechanisms by which ecdysone is increased in males versus females and whether ecdysone signaling directly inhibits EGFR function in the testis.

**Figure 4.**
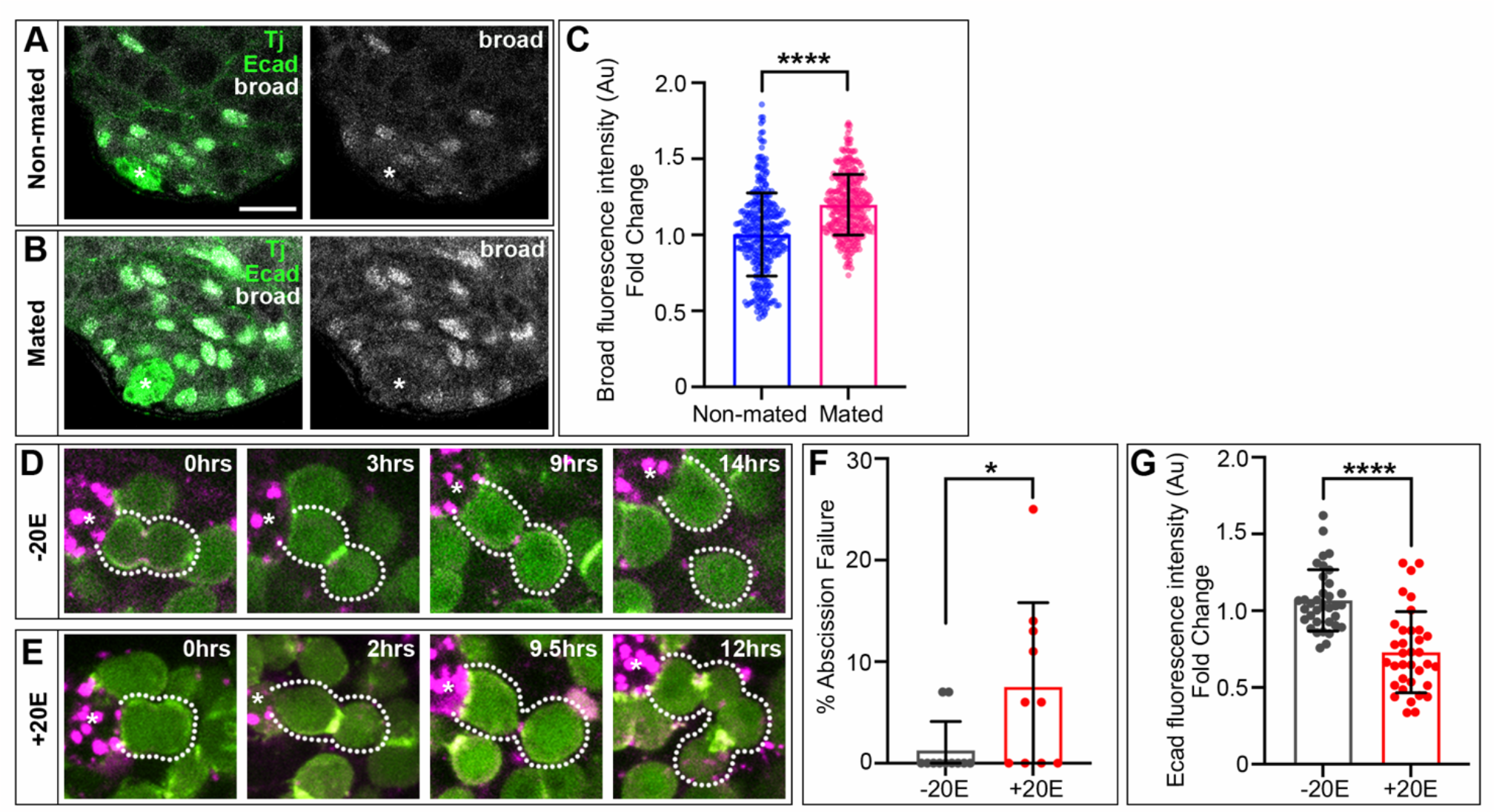
Ecdysone signaling disrupts somatic encystment, causing GSC abscission failure. (A-B) Immunostaining of Tj, E-cad, and Broad in non-mated and mated males. (C) Quantification of broad fluorescence intensity represented as fold change in testis soma. (D-E) Live imaging of nanos-ABD-moe::GFP and Myo-mCherry (sqh-sqh::mCherry) showing (D) successful completion of abscission without 20E feeding and (E) abscission failure with 20E feeding. (F) Quantification of GSC abscission failure in testes from vehicle control fed and 20E fed males. (G) Quantification of E-cad fluorescence intensity represented as fold change relative to the vehicle fed controls.

### Mating-induced GSC abscission failure is transient

In female flies, the effect of ecdysone increases on GSCs upon mating is acute and resolves as soon as sperm is lost from the seminal receptacle.^5^ To determine if the effects of mating on male GSCs are also transient, we allowed flies to rest for 1 day in the absence of females after extensive mating and performed live imaging to assess GSC cytokinesis. While not completely restored to non-mated levels, we found that GSC abscission failure was significantly reduced after only 1 day of rest (Fig.5B,E), suggesting that these defects are also transient and were ameliorated when the stress of mating was removed. This acute recovery effect on stem cell function is consistent with the transient effects of ecdysone in females^5^ and further suggests that ecdysone is the cause of GSC abscission failure in males.

**Figure 5.**
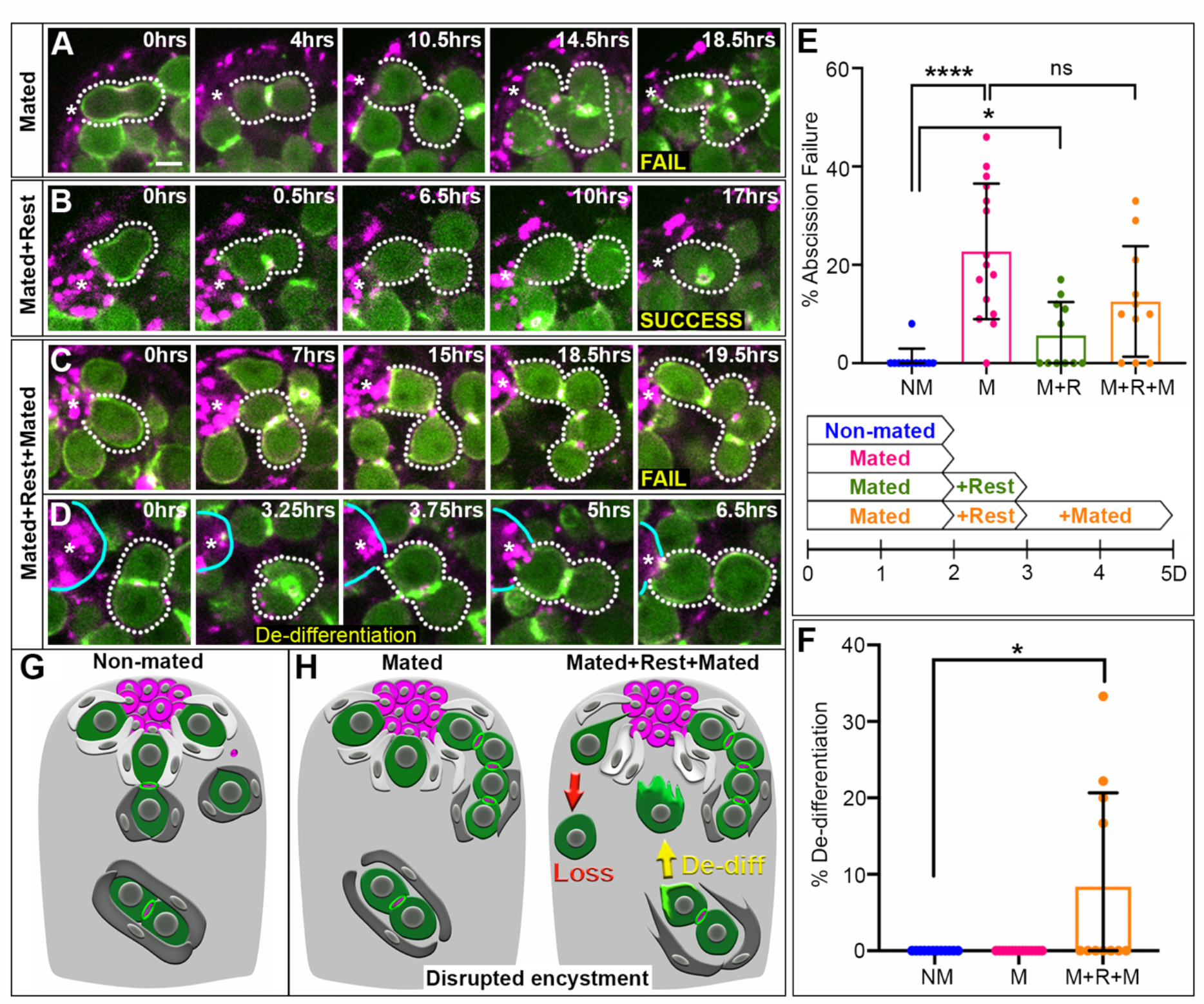
Mating-induced GSC abscission failure is transient but reoccurs with repeated bouts of mating. (A-C) Time-lapse of nanos-ABD-moe::GFP and Myo-mCherry (sqh-sqh::mCherry) during GSC divisions in testes of (A) 2 days mated (B) 2 days mated + 1 day of rest and (C) 2 days mated 1 day of rest + 2 days mated males. (D) De-differentiation of a 2-cell cyst in 2 days mated + 1 day of rest + 2 days mated testes. (D’) Diagram of a model for loss of somatic encystment enabling germ cell de-differentiation (E) Quantification of GSC abscission failure along with diagram showing mating paradigm. (F) Quantification of the percent of GSCs resulting from de-differentiation events. (G-H) Diagrams of the testis under non-mated and mated conditions. (G) Testes from non-mated males exhibit intact soma-germline contacts and completion of GSC cytokinesis. (H) Testes from mated males have disrupted soma-germline contacts, causing GSC abscission failure. Repeated bouts of a mating leads to additional defects including increased GSC loss and restoration of GSCs through de-differentiation. While still inducing GSC abscission failure, we propose a model whereby decreased somatic encystment with repeated mating facilitates germ cell de-differentiation and restoration of the GSC pool.

### Mimicking natural mating events causes abscission failure and de-differentiation to restore the stem cell pool

Intermittent bouts of mating are likely the natural experience of flies in the wild. Previous work has shown that simulating natural mating over an extended period induces GSC loss and de-differentiation of germ cells to replenish the stem cell pool.^3^ To investigate the early effects of natural mating on the GSC population, we allowed males to mate for 2 days, followed by a day of rest, and then repeated 2 days of mating. Interestingly, despite GSC abscission failures recovering with rest (Fig.5B,E), when flies underwent a second round of extensive mating, the rate of abscission failure, again, significantly increases (Fig.5C,E). Additionally, in this brief mating paradigm, we observed germ cell phenotypes normally seen with prolonged natural mating, including an increase in GSC loss and restoration of GSCs through an increase in frequency of symmetric renewal, the process of a newly produced GSC daughter rotating into the hub and becoming a new stem cell (Fig.S2). Importantly, we also observed 7 instances of de-differentiation (Fig.5D)—a behavior we have never observed in non-mated testes (Fig.5F). Together, these data suggest that even short bouts of natural mating causes GSC defects and increased GSC loss from the niche that requires induction of symmetric renewal and de-differentiation to restore the stem cell pool.

Our data supports a model in which extensive, short-term mating induces a burst of ecdysone in males, just as it does in females. However, rather than increasing GSC number as it does in the ovary, increased ecdysone signaling in the testis causes GSC abscission failure.

Furthermore, our results suggest that abscission failure results from diminished encystment mediated by decreased EGFR pathway activation. As EGFR signaling is the critical pathway for Rac1-dependent cytoskeletal changes required for germline enclosure, lowered EGFR activity upon mating impairs somatic encystment leading to decreased E-cad accumulation at soma-germline interfaces. Ultimately, defects in somatic encystment prevent robust completion of GSC abscission leading to retention of GSC daughters at the niche and failure to release germ cells to differentiate.

We have identified an immediate, detrimental effect of acute mating on the GSC population of the testis that compromises homeostatic conditions. Despite mating increasing demand for sperm, this immediate defect *decreases* release of differentiating germ cells from the niche.

What could explain this contradiction? One possibility is that enough sperm is still produced despite abscission failures to prevent selection against this acute defect in response to ecdysone. However, a more intriguing possibility is that slight decreases in encystment upon mating might be beneficial to the tissue. Previous work has shown that de-differentiating germ cells become highly protrusive as they move back to the niche—something that cannot happen if those cells were still tightly encysted by somatic cells.^22,23^ Indeed, our own live imaging reveals protrusions in de-differentiating germ cells before contacting the niche (Fig.5D), suggesting that decreases in somatic encystment, while negatively impacting GSC abscission, might serve an adaptive function to retain the stem cell pool by facilitating de-differentiation (Fig.5G-H). Importantly, persistent loss of encystment can cause over-proliferation of germ cells.^24^ Therefore, what may promote plasticity early to preserve stem cells might also explain why mated males are susceptible to developing tumors with age.^4^ Lastly, our work provides new insights into the acute effects of mating stress on the testis and reveals a role for hormonal signaling, but not germ cell loss, on the regulation of soma-germline interactions.

Given the requirement for soma-germline interactions during sperm production across species^7^, it will be interesting to determine the degree of conservation in these effects of mating on stem cell biology and germline plasticity in other systems.

## Acknowledgements

We thank Lisha Shao for the optogenetic fly stocks. A special thanks to the Von Reyn Lab for assistance with optogenetic experiments (provided RedBeam Mini LED and food vials supplemented with all trans-Retinal). We also thank Kathy Lindsley from Evident Scientific and Drexel’s Cell Imaging Center for their technical support with confocal imaging. Another thanks to Drexel undergraduate and documented dreamer, Lakshmi Parvathinathan, for her assistance with fly maintenance. Finally, thank you to Drexel graduate student, Ido Keren, for contributing intellectually to these studies. This work was supported by NIH R01 GM138705 (to KFL) and NIH 3R01GM138705-02S1 (to KFL in support of TVR).

## Author contributions

Tiffany V. Roach and Kari F. Lenhart designed experiments and wrote the manuscript. Tiffany V. Roach performed the experiments and analyzed the results.

## STAR Methods

### Key resources table

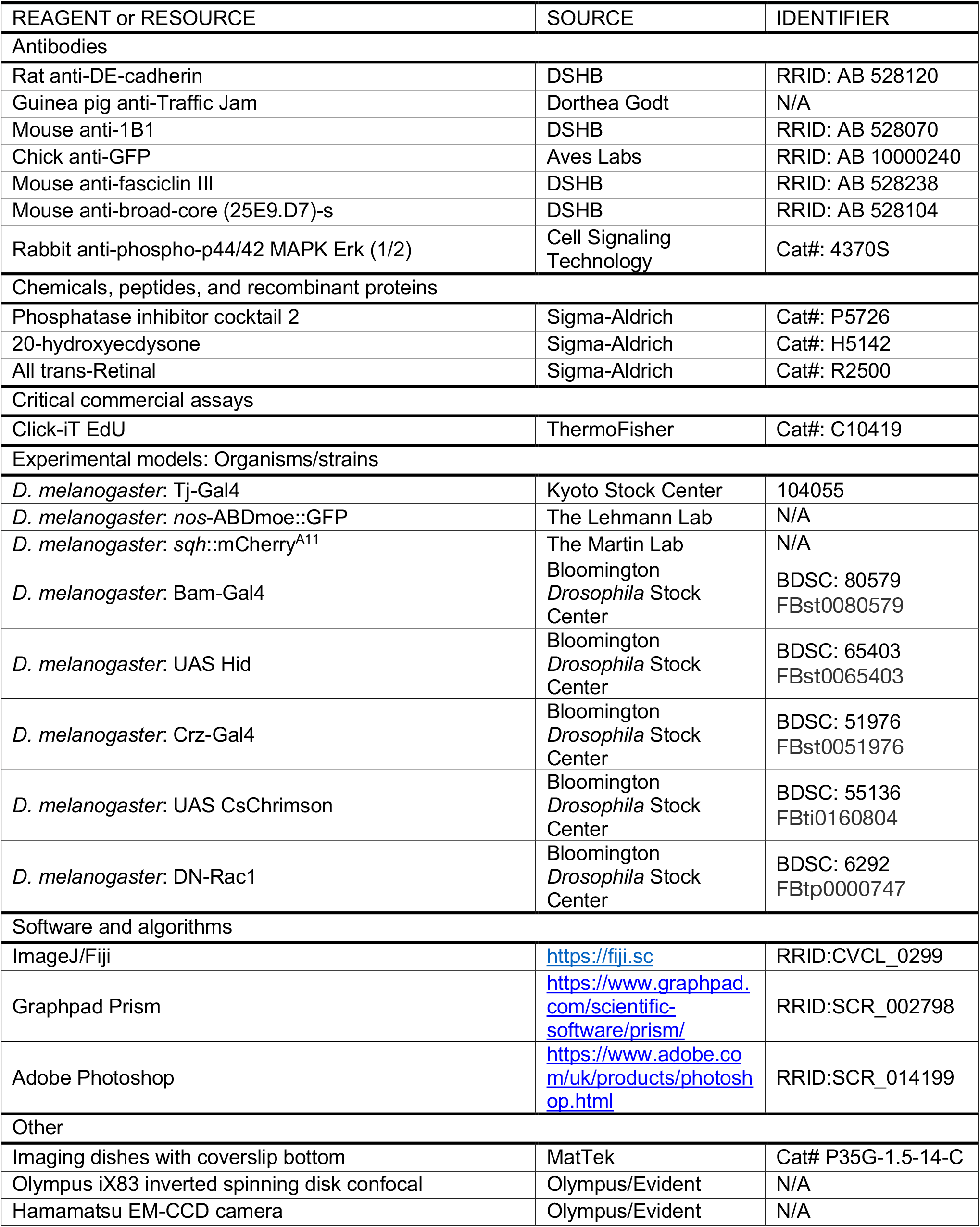
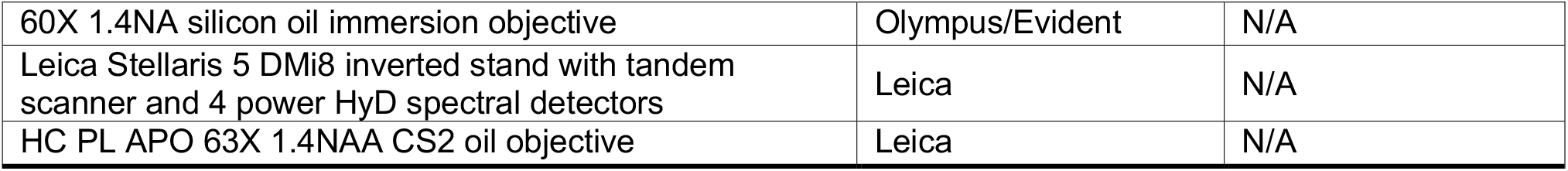

## Contact for Reagent and Resource Sharing

Requests for further information and for reagent or resource sharing should be directed to the Lead Contact Kari F. Lenhart

## Experimental Model and Subject Detail

*Drosophila melanogaster* stocks were maintained on BDSC standard cornmeal medium in vials or bottles. All crosses and males subjected to mating protocols were kept at 25° unless otherwise indicated. Fly stocks included: Traffic Jam Gal4 (Kyoto Stock Center) to drive transgene expression in somatic cells of the testis, nos-ABDmoe::GFP (Gift from Ruth Lehmann, MIT) to visualize F-actin in early stage germ cells in live imaging experiments, *sqh*::mCherry^A11^ (Gift from Adam Martin, MIT) to visualize myosin localization in live imaging experiments, Crz Gal4 (Gift from Lisa Shao; BDSC: 51976) to drive transgene expression in a subset of neurons responsible for stimulating ejaculation in the absence of mating, and UAS CsChrimson (Gift from Lisa Shao; BDSC: 55136) to sensitize neurons for optogenetic stimulation with red LED. Stocks from the Bloomington Stock Center included: UAS DN-Rac1 (BDSC: 6292) to express in somatic cells of the testis and disrupt their encystment of germ cells, Bam Gal4 (BDSC: 80579) to express an apoptotic transgene in early differentiating germ cells, and UAS Hid (BDSC: 65403) to express in early differentiating germ cells in the absence of mating.

## Methods Details

### Mating protocol

Young males underwent 2 days of mating in a 3:1 virgin female to male ratio per vial (flipping onto new food and replacing female virgins with new ones each day)—which we refer to as extensive or acute mating. Age-matched non-mated virgin males were kept together in a vial in tandem and treated identically. Male courtship was confirmed to occur throughout the full 2 days of mating. At the end of the mating paradigm, all males were sacrificed, and their testes dissected for live imaging.

### Time lapse imaging

Extended time-lapse imaging and culture conditions were adapted from (Lenhart and DiNardo, 2015; Lenhart et al., 2019; and Sheng and Matunis). Testes were dissected from 0–3-day old males in Ringers solution and mounted onto a poly-lysine-coated coverslip bottom of an imaging dish (MatTek). Ringers was removed and imaging media (15% fetal bovine serum, 0.5× penicillin/streptomycin, 0.2 mg/ml insulin in Schneider’s insect media) was added. Testes were imaged every 15 or 30 minutes for up to 24hrs on an Olympus iX83 with a Yokagawa CSU-10 spinning disk scan head, 60X 1.4NA silicon oil immersion objective, and Hamamatsu EM-CCD camera using 1 μm Z step size (40micron stacks). Experiments were repeated a minimum of two times and at least 10 testes were analyzed for each genotype/condition.

### Analysis of GSC and germ cell behaviors from live imaging

Abscission failure was defined as a GSC that entered mitosis while remaining connected to its daughter cell by a stable intercellular bridge for at least 2 consecutive time-points; whereas successful abscission was defined as an individual GSC that completely separated from its daughter cell prior to entering its next mitosis. The rate of abscission failure was calculated by dividing the total number GSC divisions with abscission failure by the total number of GSC divisions per testis. For cell cycle timing, we quantified M phase as visual production of two daughter cells (telophase or after) and the length of the cell cycle was determined for those GSCs in which we observed daughter cell production twice within the same imaging period. For quantification of either symmetric renewal or de-differentiation, the total number of GSCs acquired through each event was divided by the total number of GSCs at the end of the imaging session and reported as a percentage. For quantification of GSC loss, the total number of GSCs which lost contact with the hub were divided by the total number of GSCs at the beginning of the imaging and reported as a percentage.

### S phase labeling and analysis

To label cells in S phase, testes were dissected and incubated in 10μM of EdU for 15min prior to fixation. Testes were processed for immunohistochemistry (below), followed by visualization of the Edu label with Click-it Alexa Fluor 647 Edu kit (Invitrogen). An S phase index was determined by dividing the total number of GSCs that were Edu-positive by the total number of GSCs per testis for each condition.

### Immunostaining

Immunostaining was performed as previously described (Lenhart and DiNardo, 2015; Kari Lenhart et al., 2019; and Terry et al., 2006). In short, testes were dissected in Ringers solution and fixed for 30 minutes in 4% formaldehyde in Buffer B followed by multiple washes in PBSTx and blocking in 2% normal donkey serum. Testes were incubated in primary antibodies at 4°C for a least over one night, washed multiple times, and then incubated in appropriate secondary antibodies for 1 hour at room temperature. After additional washes, testes were equilibrated in a solution of 50% glycerol and then mounted on slides in a solution of 80% glycerol. Primary antibodies: rat anti-DE-cadherin (Developmental Studies Hybridoma Bank [DSHB], 1:20), mouse anti-1B1 (DSHB, 1:50), chick anti-GFP (1:1,000, Aves labs), guinea pig anti-traffic jam (Dorothea Godt, 1:5,000), mouse anti-fasciclin III (DSHB, 1:50), rabbit anti-phospho-p44/42 MAPK Erk (1/2) (Cell signaling technology, 1:100), and mouse anti-broad-core (25E9.D7)-s (DSHB, 1:50). Secondary antibodies: Alexa fluor-488, -Cy3, and -Cy5 (1:125) obtained from ThermoFisher Scientific. All antibodies have been previously verified by the *Drosophila* community.

Immunolabeling against dpErk was performed using the methods described above with the following exceptions. Washes were done using PBS supplemented with phosphatase inhibitor cocktail 2 (1:100, P5726, Sigma-Aldrich). This mixture was used in place of Buffer B for the 30min fixation. Dissections were conducted on ice and each incubation/wash step was done at 4°C.

### Quantification of fluorescence intensities

For analysis of E-cadherin accumulation, we traced the soma-germline interfaces of 2-cell cysts using freehand line tool with a pixel width of 3 in ImageJ. Additionally, 3 niche-niche interfaces were measured and averaged as an internal reference as well as 3 background traces to subtract. Thus, the following formula was used: (mean 2-cell cyst trace – mean average background) / (mean average reference – mean average background). The average mean for all 2-cell cysts from the non-mated testes was determined and used to normalize each value from both non-mated and mated testes to reveal fold change differences. For each genotype, a minimum of 10 testes and 40 2-cell cysts were analyzed were analyzed across at least two independent trials.

For both dpErk and Broad analyses, ROIs were drawn around somatic nuclei as defined by Traffic jam accumulation and then mean fluorescence intensity of dpErk and Broad were measured separately in ImageJ. Again, the average mean for all Tj+ cells within 50 microns from the testis tip in the non-mated testes was determined and used to normalize each value from both non-mated and mated testes to reveal fold change differences. All experiments and respective controls were done in tandem and imaged with the same acquisition settings. For each genotype, a minimum of 6 testes and 345 nuclei were analyzed across at least two independent trials.

All images of fixed and immunostained testes were acquired using a Leica Stellaris 5 DMi8 inverted stand with tandem scanner; 4 power HyD spectral detectors; and HC PL APO 63x/1.4NA CS2 oil objective using LAS X software.

### Optogenetic Activation

The parental cross (UAS CsChrimson x Crz Gal4) was reared on food supplemented with 1:500 trans retinal in ethanol in the dark at 25°C. Newly eclosed virgin males carrying the UAS CsChrimson and Crz Gal4 were collected onto food containing 1:250 all trans-Retinal (R2500, Sigma-Aldrich) for 3 days. Then, while in the dark, males were anesthetized on ice, transferred to empty vials, and allowed 5min to sufficiently wake. Next, males were exposed to red LED light (∼650nm, RedBeam™ Mini) for 3, 15min activations with at least 45min of rest in their respective food vials in between exposures (Zer-Krispil et al., 2018). We visually confirmed successful activation of neurons and resulting ejaculation with this paradigm. For experiments, the light was positioned to shine directly into the vial containing males and wrapped in foil to restrict ambient light. Additionally, this apparatus was placed in a black box at room temperature for the duration of the experiments. This procedure was repeated for 2 consecutive days to closely mimic the mating experience of our acute mating paradigm and males were subsequently sacrificed for live imaging.

### Ecdysone Feeding Experiments

Young virgin males were fed a mixture of 20-hydroxyecdysone (1mM, Sigma-Aldrich Cat. No. H5142), 1% ethanol, 100% Organic Apple Juice (Nature’s Promise), and blue dye (McCormick) on filter paper. Half the males were fed the same mixture but lacking 20E. The food vials, filter papers, and mixtures were replaced each day for 2 days to match the timeframe of the acute mating protocol. Following this, testes were dissected for live imaging.

### Quantification, Statistical Analysis, and Image processing

Time-lapse images were analyzed and Z projections generated using ImageJ software. All graphical representations of data and statistical analysis were performed in Graphpad Prism (non-parametric Mann Whitney U test). Error bars represent standard deviation. *n* and p values indicated in figure legends. Figures were generated using BioRender.com and Adobe Photoshop.

