## Supplemental Figures for "Mating-induced ecdysone in the testis disrupts soma-germline contacts and stem cell cytokinesis"

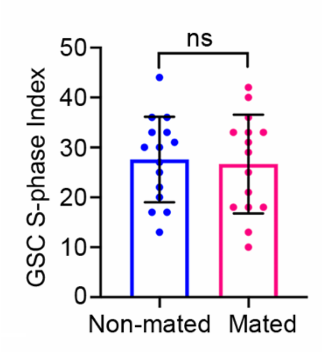


**Supplemental Figure 1.** S phase index of GSCs in non-mated vs mated males. A minimum of 14 testes were analyzed per condition.
How d

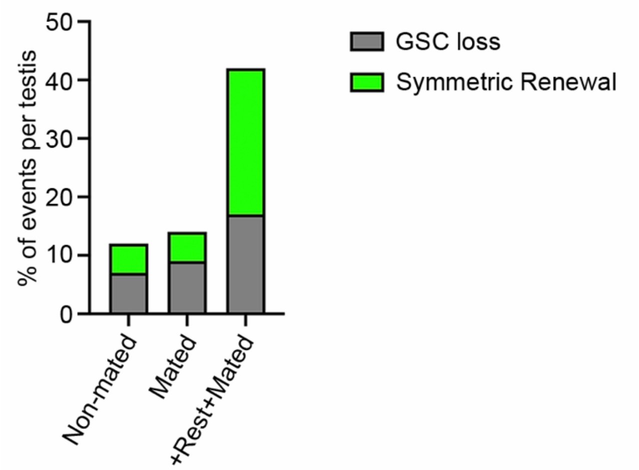


**Supplemental Figure 2.** Number of events for GSC loss and symmetric renewal in testes from non-mated, mated, and +Rest+Mated males. A minimum of 11 testes were analyzed for quantification of GSC loss and symmetric renewal events.
